# TorsinB overexpression prevents abnormal twisting in DYT1 dystonia mouse models

**DOI:** 10.1101/836536

**Authors:** Jay Li, Chun-Chi Liang, Samuel S. Pappas, William T. Dauer

## Abstract

Genetic redundancy can be exploited to identify therapeutic targets for inherited disorders. An example is DYT1 dystonia, a neurodevelopmental movement disorder caused by a loss-of-function (LOF) mutation in the *TOR1A* gene encoding torsinA. Prior work demonstrates that torsinA and its paralog torsinB have conserved functions at the nuclear envelope. This work established that low neuronal levels of torsinB dictate the neuronal selective phenotype of nuclear membrane budding. Here, we examined whether torsinB expression levels impact the onset or severity of abnormal movements, or neuropathological features in DYT1 mouse models. We demonstrate that torsinB levels bidirectionally regulate these phenotypes. Reducing torsinB levels causes a dosedependent worsening whereas torsinB overexpression rescues torsinA LOF-mediated abnormal movements and neurodegeneration. These findings identify torsinB as a potent modifier of torsinA LOF phenotypes and suggest that augmentation of torsinB expression level may retard or prevent symptom development in DYT1 dystonia.

## Introduction

DYT1 dystonia is a dominantly inherited movement disorder that is caused by 3-bp in-frame deletion (ΔE mutation) in the *TOR1A* gene that encodes the torsinA protein [1]. Only ~30% of mutation carriers exhibit symptoms, which vary in severity from mild to severely debilitating [2, 3]. Treatments include deep brain stimulation, which is invasive, and anticholinergic drugs, which provide incomplete relief and are plagued by side effects [4, 5]. These empiric treatments suppress symptoms; no therapies are based on disease pathogenesis or alter the emergence of symptoms.

TorsinA is a nuclear envelope/endoplasmic reticulum (NE/ER) resident AAA+ protein (<u>A</u>TPase <u>A</u>ssociated with diverse cellular <u>A</u>ctivities) [6]. Multiple lines of evidence demonstrate that the DYT1 mutation impairs torsinA function [7–10]. The DYT1 mutation reduces protein stability and impairs interaction with cofactors (LAP1 and LULL1) that appear important for torsinA ATPase activity [10–12].

Prior work demonstrates conserved functions for torsinA and torsinB. Both proteins are expressed ubiquitously [13]. Their sequences are 68% identical and 85% similar, and they share cofactors LAP1 and LULL1 [14–16]. TorsinA null mice and mice homozygous for the DYT1 mutation exhibit neural-selective abnormalities of NE structure (NE “budding”) [11, 17]. Several observations suggest that this neural specificity results from markedly lower levels of torsinB in neuronal compared with non-neuronal cells. The appearance of neuronal NE budding in torsinA mutants coincides with lower levels of torsinB during early brain maturation [18]. shRNA knockdown of torsinB in torsinA null non-neuronal cells recapitulates the “neuronal” NE phenotype [17]. Moreover, conditional CNS deletion of both torsinA and torsinB causes NE budding in neuronal and non-neuronal (e.g. glia) cells, and overexpressing torsinB significantly reduces NE budding in torsinA null developing neurons *in vitro* [18].

Based on these data, we hypothesized that altering torsinB levels would bidirectionally modulate motor and neuropathological phenotypes of DYT1 mouse models. We pursued epistatic analyses of torsinA loss-of-function (LOF) and both torsinB reduction and torsinB overexpression, assessing established torsinA-related neuropathological and behavioral phenotypes. We demonstrate that torsinB bidirectionally modifies these phenotypes. Reducing levels of torsinB in DYT1 models (*Tor1a*^-/-^ or *Tor1a*^ΔE/-^) causes a dose-dependent worsening of behavioral and neuropathological phenotypes. Conversely, overexpressing torsinB from the ROSA26 locus dramatically reduced the emergence of motor and neuropathological phenotypes in two DYT1 models. Our findings demonstrate that torsinB is a genetic modifier of torsinA LOF disease-related phenotypes and suggest that enhancing torsinB function may be a viable therapeutic strategy in DYT1 dystonia.

## Results

### TorsinB null mice exhibit no apparent organismal or neuropathological phenotypes

To study the role of torsinB as a modifier of torsinA LOF, we first assessed whether torsinB null mice exhibit any pathological phenotypes. Consistent with previous studies, intercrosses of *Tor1b*^+/-^ mice yielded *Tor1b^-/-^* mice that were indistinguishable from littermate controls [18]. *Tor1b*^-/-^ mice gain weight similarly to littermate controls (Figure S1A), and do not exhibit brain abnormalities when assessed by Nissl stain (Figure S1B). Glial fibrillary acidic protein (GFAP) immunohistochemistry did not demonstrate any areas of gliosis (Figure S1C). *Tor1b*^-/-^ mice exhibit normal cortical thickness (Figure S1D) and do not display abnormal limb clasping during tail suspension (data not shown). These findings enabled us to assess the role of torsinB levels on torsinA LOF-related phenotypes without the confound of additive abnormalities.

### TorsinB deletion worsens torsinA-related motor and neuropathological phenotypes

Emx1-Cre expresses in the majority of cortical and hippocampal neurons [19]. This Cre field includes the motor cortex, which is a critical component of the forebrain cortico-striatal network thought to participate in the expression of dystonic symptoms [20–22]. Disruption of this network has been observed in several dystonia mouse models [23–26]. *Emx1-Cre* conditional deletion of torsinA (Emx1-CKO, Table S1, Figure 1A) reduces cortical thickness (Figure 1D), but does not significantly alter the number of CUX1+ (marker for cortical layer II-IV) or CTIP2+ (marker for cortical layer V-VI) cortical neurons (Figure 1E-F) [27, 28]. Emx1-CKO mice appear grossly normal, and only a subset of these mice exhibit limb clasping during tail suspension (Table S1, Figure 1G) [8]. The phenotype of Emx1-CKO mice therefore avoids ceiling effects, making it an appropriate model to assess phenotypic worsening with combined torsinA and torsinB LOF. Consistent with prior studies, conditionally deleting both torsinA and torsinB with *Emx1-Cre* (Emx1-dCKO [“d” for double], Figure 1A) does not cause overt brain structural abnormalities at birth [11]: cortical thickness and the number of CTIP2+ cells do not differ significantly from littermate controls at postnatal day 0 (P0) (Figure S2A-B). At P28, however, Emx1-dCKO mice exhibit significant cell loss (Figure 1B) and profound gliosis (Figure 1C) in Cre expressing regions (cerebral cortex and hippocampus). The hindbrain, which is not in the Cre field, appears normal (Figure S2C). The cerebral cortex is significantly thinner in Emx1-dCKO than in Emx1-CKO mice (64.8% vs. 10.4% reduction compared to littermate controls, respectively; Figure 1C-D). Whereas cell counts are normal in Emx1-CKO mice, Emx1-dCKO mice exhibit significant reductions of CUX1+ and CTIP2+ neurons in sensorimotor cortex (Figure 1E-F). Consistent with the enhanced neuropathological phenotype, torsinB removal significantly worsens the behavioral phenotype (Figure 1G). Considered together with the observation that *Tor1b*^-/-^ mice appear normal, these data suggest that torsinB expression is essential on a torsinA LOF background.

**Figure 1.**
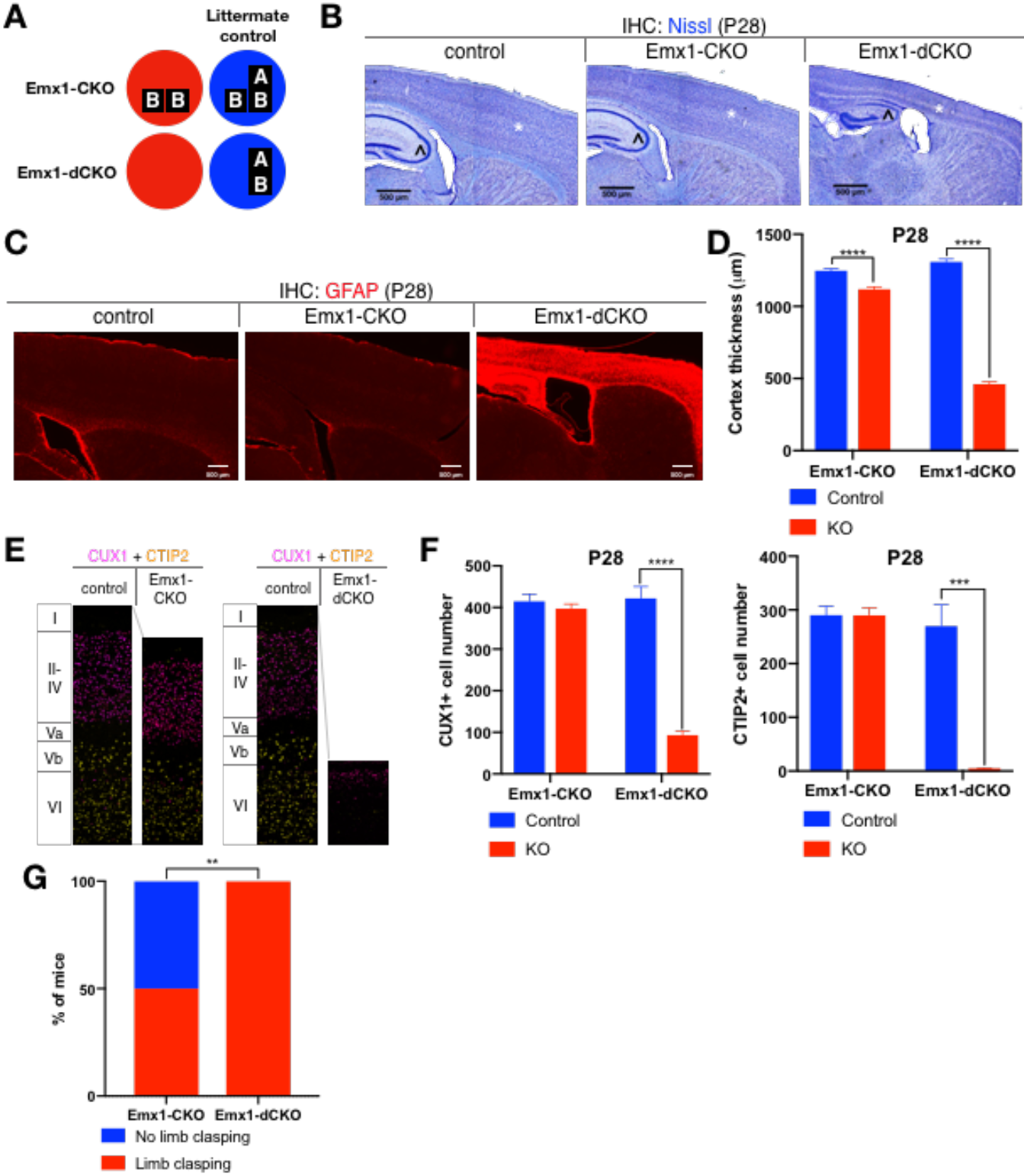
TorsinB deletion worsens torsinA-related motor and neuropathological phenotypes. (A) Illustration of examined genotypes. Each row of boxes within the circles denote the presence or absence of *Tor1a* (top row) or *Tor1b* (bottom row) alleles following Cre recombination. The presence of a letter (“A” or “B”) indicates an intact allele, whereas the absence of a letter indicates a deleted allele. (B) Nissl staining of P28 Emx1-CKO and Emx1-dCKO mice. Emx1-dCKO mice exhibit significant atrophy of *Cre*-expressing areas including cortex (*) and hippocampus (^). (C) GFAP staining of P28 Emx1-CKO and Emx1-dCKO brains. Emx1-dCKO mice exhibit severe reactive gliosis in *Cre*-expressing regions, including the cerebral cortex and the hippocampus. (D) Cortical thickness of P28 Emx1-CKO and Emx1-dCKO mice. Cortical thickness is reduced by 10.4% in Emx1-CKO mice (unpaired t-test t_16_ = 5.834, p < 0.0001; control n = 9, Emx1-CKO n = 9). Cortical thickness is reduced by 64.8% in Emx1-dCKO (unpaired t-test t_11_ = 30.16, p < 0.0001; control n = 4, Emx1-dCKO n = 9). (E) Representative images from CUX1 and CTIP2 stained cerebral cortex of Emx1-CKO and Emx1-dCKO mice and their respective littermate controls. (F) CUX1 and CTIP2 counts in P28 Emx1-CKO and Emx1-dCKO mice. CUX1+ neurons are not significantly reduced in Emx1-CKO mice (unpaired t-test t_16_ = 0.8469, p = 0.4095; control n = 9, Emx1-CKO n = 9). CUX1+ cells are significantly reduced in Emx1-dCKO mice (77.0% reduction; unpaired t-test t_8_ = 12.86, p < 0.0001, control n = 4, Emx1-dCKO n = 6). CTIP2+ neurons are not significantly reduced in Emx1-CKO sensorimotor cortex (unpaired t-test t_16_ = 0.02552, p = 0.98; control n = 9; Emx1-CKO n = 9). CTIP2+ neurons are significantly reduced in Emx1-dCKO sensorimotor cortex (98.6%; unpaired t-test t_7_ = 7.636, p = 0.0001, control n = 4, Emx1-dCKO n = 5). (G) Proportion of Emx1-CKO and Emx1-dCKO mice exhibiting limb clasping during tail suspension. A significantly greater proportion of Emx1-dCKO compared to Emx1-CKO mice exhibit limb clasping during tail suspension (Chi square test χ^2^ = 9.1, p = 0.0026; Emx1-CKO n = 12, Emx1-dCKO n = 14).

### TorsinB deletion causes a dose-dependent worsening in the *Tor1a*^ΔE/-^ Emx1-SKI mouse model

To investigate the impact of torsinB reduction in the presence of DYT1 (ΔE) mutant torsinA, we examined ΔE torsinA “selective-knock-in” (SKI) mice [8]. In Emx1-SKI mice (*Emx1-Cre*; *Tor1a^ΔE/flx^*), a floxed copy of torsinA is deleted upon *Cre* recombination, leaving isolated expression of ΔE disease mutant torsinA (*Tor1a*^ΔE/-^) in *Emx1-Cre* expressing excitatory forebrain neurons (Table S1, Figure 2A). We performed a torsinB gene dosage study on Emx1-SKI mice assessing mutants heterozygous (Emx1-SKI-B1) or homozygous (Emx1-SKI-B0) for a *Tor1b* null allele (Figure 2A, S3A).

**Figure 2.**
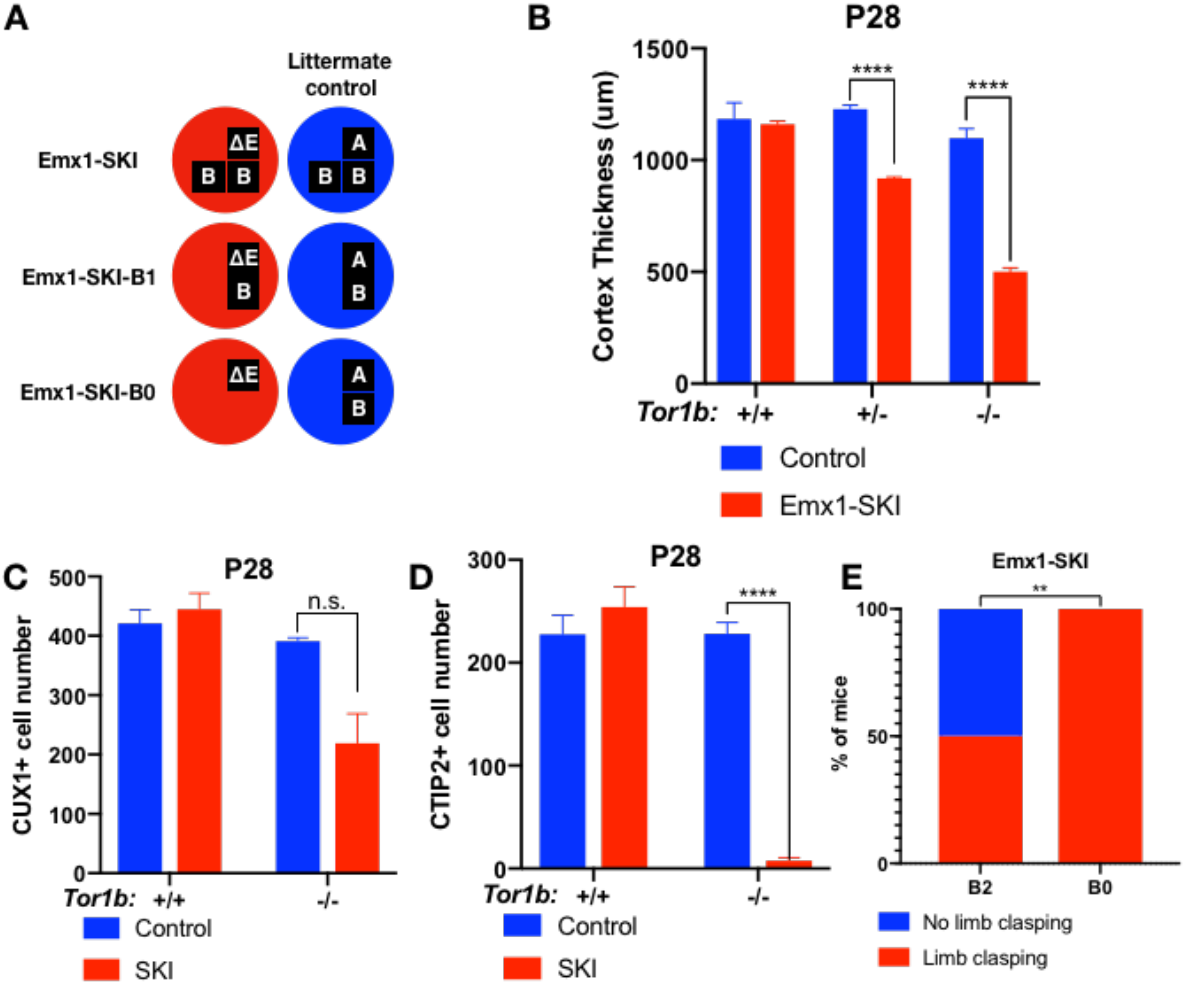
TorsinB deletion causes a dose-dependent worsening in the *Tor1a*^ΔE/-^ Emx1-SKI mouse model. (A) Illustration of examined genotypes. Each row of boxes within the circles denote the presence or absence of *Tor1a* (top row) or *Tor1b* (bottom row) alleles following Cre recombination. The presence of a letter (“A”“ΔE” or “B”) indicates an intact allele, whereas the absence of a letter indicates a deleted allele. (B) Cortical thickness of P28 Emx1-SKI, Emx1-SKI-B1, and Emx1-SKI-B0 mice. TorsinB loss dose dependently reduces cortical thickness in Emx1-SKI mice in a dose-dependent manner (2% reduction in Emx1-SKI, 25.2% reduction in Emx1-SKI-B1, and 54.5% decrease in Emx1-SKI-B0; two way ANOVA main effect of background genotype F_1,13_ = 128. 3, p < 0.0001, age F_2,13_ = 62.65, p < 0.0001, and interaction F_2,13_ = 36.98, p < 0.0001; Sidak’s multiple comparisons test p = 0.9323 for *Tor1b^+/+^*, p < 0.0001 for *Tor1b^+/-^*, P < 0.0001 for Tor1b^-/-^; *Tor1b^+/+^* control n = 3; Emx1-SKI n = 5, *Tor1b^+/-^* control n = 3, Emx1-SKI n = 3, *Tor1b^-/-^* control n = 2, Emx1-SKI n = 3). (C) CUX1+ cell counts in sensorimotor cortex of P28 Emx1-SKI and Emx1-SKI-B0 mice. There is no significant reduction in the number of CUX1+ cells in Emx1-SKI sensorimotor cortex (unpaired t-test t_16_ = 0.8469, p = 0.4095; control n = 3, SKI n = 5). Simultaneous deletion of two torsinB alleles reduces CUX1+ cell counts by 44.0% (unpaired t-test t_3_ = 2.655, p = 0.0766; control n = 2, SKI n = 3) though this reduction does not reach statistical significance. (D) CTIP2+ cell counts in sensorimotor cortex of P28 Emx1-SKI and Emx1-SKI-B0 mice. CTIP2+ neuronal cell counts are not reduced in Emx1-SKI mice (unpaired t-test t_6_ = 0.8844, p = 0.4105; control n = 3, SKI n = 5). CTIP2+ cell counts in Emx1-SKI-B0 are significantly reduced by 96.6% (unpaired t-test t_3_ = 24.44, p <0.0001; control n = 2, SKI n = 3). (E) Proportion of Emx1-SKI and Emx1-SKI-B1 mice exhibiting limb clasping during tail suspension. A significantly greater proportion of Emx1-SKI-B1 mice exhibit limb clasping during tail suspension compared to Emx1-SKI (Chi square test χ^2^ = 7.441, p = 0.0064; Emx1-SKI n = 12, Emx1-SKI-B1 n = 11).

The cortical thickness of Emx1-SKI mice does not differ significantly from littermate controls and these mice have normal numbers of CUX1+ and CTIP2+ cells (Figure 2B-D). Ablation of a single torsinB allele (Emx1-SKI-B1) significantly reduces cortical thickness but not CUX1+ or CTIP2+ neuron counts (Figure 2B, S3C). The cortical thickness of mice with complete loss of torsinB (Emx1-SKI-B0) is normal at birth (Figure S3B), but by P28 is dramatically reduced to an extent further than in Emx1-SKI-B1 mice (Figure 2B). At P28, Emx1-SKI-B0 mice also exhibit a nonsignificant reduction of CUX1+ neurons (60.9%, p = 0.0766) and a significant reduction of CTIP2+ neurons (Figure 2C-D). These data highlight two points: 1) torsinB reduction worsens phenotypes when ΔE torsinA is present, and 2) this worsening occurs in a dose-dependent manner. Consistent with a dose-dependent worsening of neuropathology, removal of a single torsinB allele (Emx1-SKI-B1) significantly worsened the behavioral phenotype (Figure 2E).

### A novel Cre-dependent torsinB overexpression allele

Having established that reducing torsinB levels exacerbates multiple torsinA mutant neuropathological and behavioral phenotypes, we explored whether enhancing torsinB levels could suppress or prevent these same features. We generated a torsinB overexpression (B-OE) allele by knocking the *Tor1b* cDNA into the ROSA26 locus (Figure 3A). The *Tor1b* cDNA is preceded by a floxed “stop” cassette, rendering this allele Cre dependent. This design allows for Cre activation of torsinB overexpression in the same spatiotemporal pattern as Cre deletion of the floxed *Tor1a* allele.

**Figure 3.**
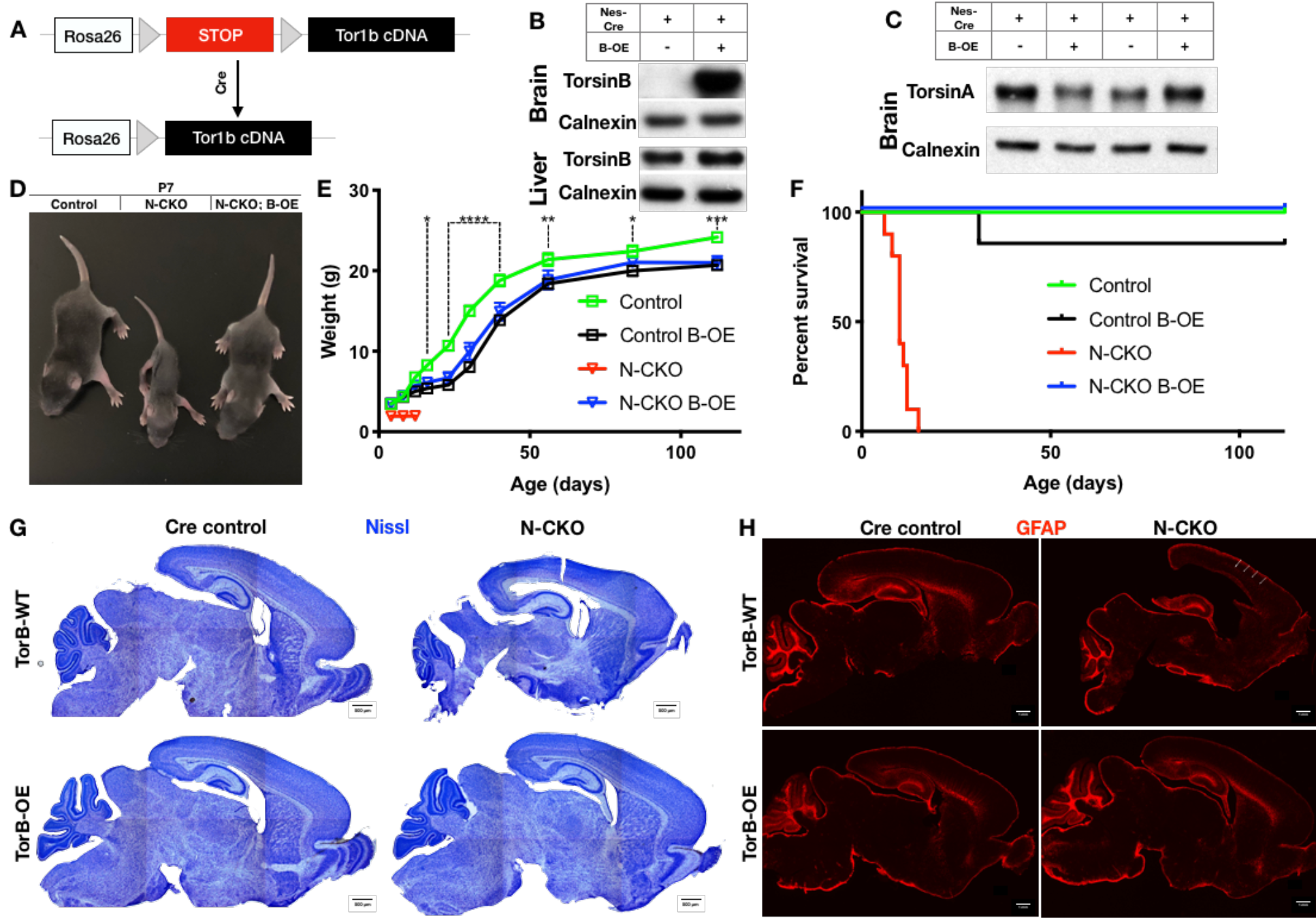
TorsinB augmentation prevents torsinA-related lethality and neuropathology. (A) Cartoon illustration of ROSA26 locus engineered to express torsinB. Top: In absence of *Cre*, torsinB expression is prevented by a floxed “STOP” cassette. Bottom: TorsinB is expressed following *Cre* deletion of the floxed “STOP” cassette. (B) Western blot analysis of whole brain and liver lysates probed with an anti-torsinB antibody. Mice expressing both the *Nestin-Cre* and B-OE alleles exhibit cre-dependent overexpression of torsinB. (C) Western blot analysis of whole brain lysates probed with anti-torsinA antibody. TorsinB overexpression does not significantly alter torsinA expression. (D) Growth curves of *Cre* control, *Cre* control;B-OE, N-CKO, and N-CKO;B-OE mice. TorsinB overexpression almost entirely restores growth in N-CKO mice (two-way repeated measures ANOVA main effect of genotype F_2,17_ = 14.16, p = 0.0002, main effect of age F_9,153_ = 775.7 p < 0.0001, interaction F_18,153_ = 6.727, p < 0.0001; asterisks denote the following p-values from Tukey’s multiple comparisons tests: * = p < 0.05, ** = p < 0.01, ** = p < 0.001, **** = p < 0.0001; *Cre* control n = 7, *Cre* control;B-OE n = 6, N-CKO;B-OE n = 7). (E) Survival curves of *Cre* control, *Cre* control;B-OE, N-CKO, and N-CKO;B-OE mice. TorsinB overexpression eliminates lethality in N-CKO mice (Gehan-Breslow-Wilcoxon method χ^2^ = 36.16, p < 0.0001; *Cre* control n = 7, *Cre* control;B-OE n = 7, N-CKO n = 10, N-CKO;B-OE n = 7). (F) Nissl staining of P8 brains from *Cre* control, *Cre* control;B-OE, N-CKO, and N-CKO;B-OE mice. TorsinB overexpression eliminates the morphological defects characteristic of N-CKO mice. (G) GFAP staining of P8 brains from *Cre* control, *Cre* control;B-OE, N-CKO, and N-CKO;B-OE mice. GFAP immunostaining illustrates the characteristic gliotic changes in N-CKO cortex (arrows), thalamus, deep cerebellar nuclei, and hindbrain (circles). TorsinB overexpression eliminates gliotic changes in N-CKO mice.

We validated the B-OE mouse line by measuring levels of torsinB in brain lysates from *Nestin-Cre* mice, where the *Cre* is active in the cells that give rise to the entire nervous system. As expected, torsinB is selectively overexpressed only in mice harboring both the *Cre* and B-OE alleles (Figure 3B). This analysis demonstrates that the B-OE allele supports marked overexpression of torsinB, likely in part because of the included woodchuck hepatitis virus posttranscriptional regulatory element. *Nestin-Cre*;B-OE mice exhibit normal torsinB expression in liver (a Cre negative tissue), demonstrating that overexpression is selective and Cre-dependent (Figure 3B). We next tested whether torsinB level impacts torsinA expression, as past studies suggest that torsinA and torsinB have reciprocal timing and tissue patterns of expression [17, 18]. TorsinB overexpression did not significantly alter torsinA expression in brain lysates, indicating that there is not a direct regulatory relationship between levels of these proteins (Figure 3C, S4A). These findings confirm that the new mouse line overexpresses torsinB in a Cre-dependent manner. We next employed this line to explore whether torsinB overexpression can rescue torsinA LOF phenotypes.

### TorsinB augmentation prevents torsinA-related lethality and eliminates neuropathological and behavioral phenotypes in N-CKO mice

Prior work demonstrates that torsinB overexpression significantly suppresses a torsinA LOF cellular phenotype *in vitro*. To determine if torsinB overexpression can rescue torsinA LOF behavioral and neuropathological phenotypes *in vivo*, we ingressed the *Cre*-dependent B-OE allele onto the *Nestin-Cre* torsinA conditional knockout (N-CKO) background. Nestin-*Cre* is active in neural progenitor cells that give rise to the entire nervous system. N-CKO mice exhibit early lethality, lack of postnatal weight gain, overtly abnormal twisting movements, and gliosis in multiple sensorimotor brain regions (Table S1) [8], robust phenotypes that can be harnessed to determine if torsinB compensates for torsinA LOF. We analyzed four genotypes: *Nestin-Cre*; *Tor1a*^flx/+^ (Cre control), *Nestin-Cre*; *Tor1a*^flx/+^; B-OE (*Cre* control;B-OE), *Nestin-Cre*; *Tor1a*^flx/-^(N-CKO), and *Nestin-Cre*; *Tor1a*^flx/-^; B-OE (N-CKO;B-OE). Consistent with prior work, N-CKO mice weighed less than their littermates controls and exhibited 100% lethality by P15 (Figure 3D-F) [8]. Strikingly, torsinB overexpression eliminated N-CKO lethality and significantly rescued postnatal growth (Figure 3D-F). As described previously, N-CKO mice demonstrated overt abnormal twisting movements and stiff postures [8]. In contrast torsinB overexpression completely eliminated these abnormal movements (data not shown). The widespread high, non-physiologic levels of torsinB produced by the B-OE allele impaired bodyweight in *Nestin*-*Cre* control mice (Figure 3E; comparing *Cre* control and *Cre* control;B-OE), but did not induce overt motor abnormalities, or reduce survival (Gehan-Breslow-Wilcoxon method χ^2^ = 1, p = 0.3173; *Cre* control n = 7, *Cre* control;B-OE n = 7). Nissl staining at P8 (prior to lethality) demonstrated the expected N-CKO neuropathology. N-CKO brains exhibited enlarged lateral ventricles and a reduction in overall brain size compared to *Cre* controls (Figure 3G, S4B). Brain morphology and size appeared normal in both *Cre* control;B-OE and N-CKO;B-OE mice (Figure 3G, S4B). Consistent with prior work, the brains of N-CKO mice exhibited GFAP staining reflecting gliosis in the cerebral cortex, thalamus, brainstem, and deep cerebellar nuclei (Figure 3H) [8]. In striking contrast, torsinB overexpression prevented these gliotic changes (Figure 3H).

### TorsinB overexpression prevents ChI degeneration and dystonic-like movements

Conditional deletion of torsinA from the forebrain using *Dlx5/6-Cre* (“Dlx-CKO”) produces a robust DYT1 model with high face and predictive validity (Table S1) [29]. These mutants develop abnormal limb and trunk twisting movements as juveniles that are responsive to anti-muscarinic drugs, mimicking key features of the human phenotype. Coincident with the emergence of these movements, these animals exhibit a highly selective loss of dorsal striatal cholinergic interneurons (ChIs), cells that have been implicated in dystonia pathophysiology in electrophysiological studies [29–31]. The disease relevant time course of motor features, clear endpoints, and predictive validity of this model provides a critical system for evaluating the effects of torsinB overexpression in a torsinA LOF pathophysiological *in vivo* setting. To explore the ability of the B-OE allele to suppress or prevent phenotypes in this model, we first confirmed that this allele functions as expected with the *Dlx5/6-Cre* transgene and assessed for any behavioral effects caused by overexpressing torsinB in this Cre field. Mice co-expressing *Dlx5/6-Cre* and the B-OE allele selectively overexpressed torsinB in the forebrain in a *Cre*-dependent manner (Figure S5A). TorsinB overexpression in the *Dlx5/6*-*Cre* field (*Dlx5/6*-*Cre*;B-OE) caused no abnormalities of postnatal growth (Figure S5B) or survival (all mice survived, data not shown) compared to mice only expressing *Dlx5/6-Cre*. Further, *Dlx5/6-Cre*;B-OE mice exhibited normal juvenile reflexes, including negative geotaxis, righting, and forelimb hang (Figure S5C).

Having confirmed the proper functioning and lack of toxicity of the B-OE allele in the *Dlx5/6-Cre* field, we examined the effect of torsinB overexpression on Dlx-CKO motor and neuropathological phenotypes. We examined four genotypes: *Dlx5/6-Cre; Tor1a^flx/+^* (Cre control), *Dlx5/6-Cre; Tor1a^flx/+^*; B-OE (Cre control;B-OE), *Dlx5/6-Cre; Tor1a^flx/-^* (Dlx-CKO), and *Dlx5/6-Cre; Tor1a*^flx/-^; B-OE (Dlx-CKO;B-OE). Overexpression of torsinB significantly suppressed abnormal twisting movements in Dlx-CKO mice. We recorded 1-minute tail suspension videos at age P70, and 2 observers blind to genotype scored each video for duration of abnormal movements. Dlx-CKO mice exhibited limb clasping for an average of 36.9 ± 5.3 s (mean ± SEM). TorsinB overexpression (Dlx-CKO;B-OE) significantly reduced the time spent clasping (2.5 ± 1.1 s, mean ± SEM) by over 10-fold (Figure 4A). Notably, the duration of clasping in Dlx-CKO;B-OE did not differ significantly from their littermate controls. We also assessed the mice for the presence of trunk twisting during tail suspension. This phenotype occurs in most (9 of 11) Dlx-CKO mice but was completely eliminated by the B-OE allele (0/15 trunk twisting, p < 0.0001 compared to Dlx-CKO, Figure 4B). Considered together, these data establish the ability of torsinB to suppress DYT1-related motor phenotypes.

**Figure 4.**
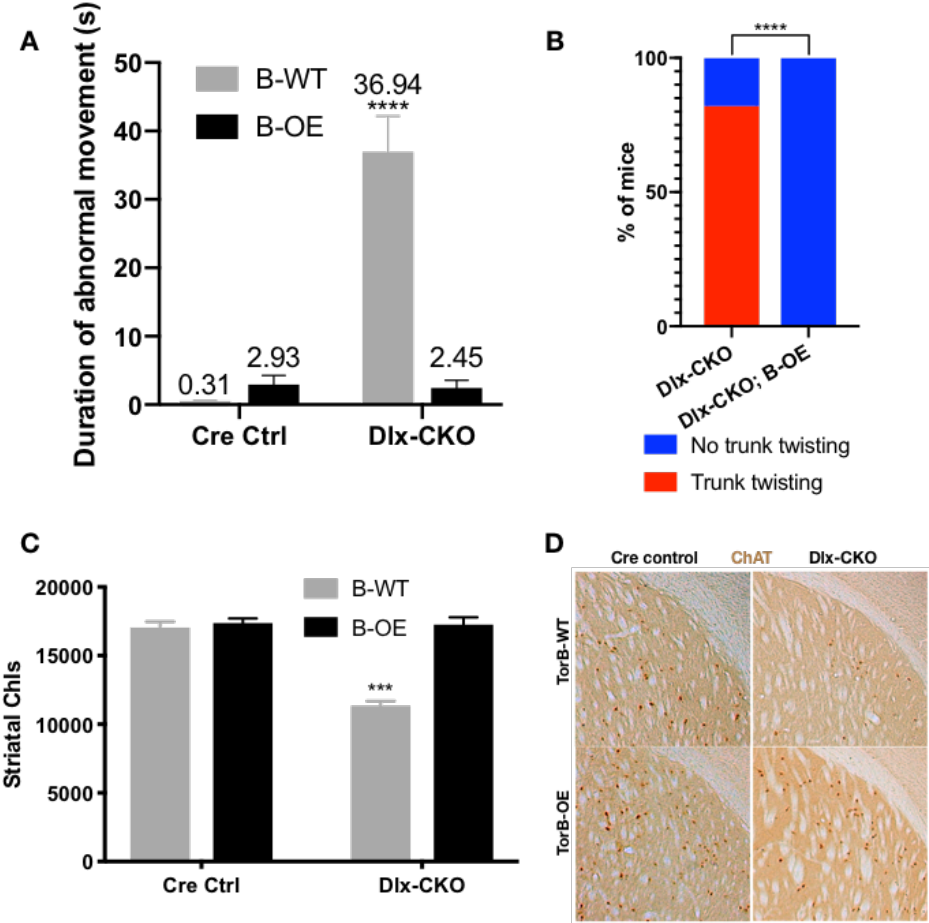
TorsinB overexpression prevents ChI degeneration and dystonic-like movements. (A) Duration of abnormal movements during one minute of tail suspension in P70 *Cre* control, *Cre* control;B-OE, Dlx-CKO, and Dlx-CKO;B-OE mice. TorsinB overexpression significantly reduces severity of limb clasping (two-way ANOVA main effect of genotype F_1,44_ = 47.45, p < 0.0001, torsinB level F_1,44_ = 36.86, p < 0.0001, and interaction F_1,44_ = 49.96, p < 0.0001; *Cre* control n = 10, *Cre* control;B-OE n = 12, Dlx-CKO n = 11, Dlx-CKO;B-OE n = 15). (B) Prevalence of trunk twisting in Dlx-CKO and Dlx-CKO;B-OE mice. TorsinB overexpression significantly reduces prevalence of trunk twisting (Chi square test χ^2^ = 18.77, p < 0.0001; Dlx-CKO n = 11, Dlx-CKO; B-OE n = 15). (C) Stereologic counts of striatal cholinergic interneurons in P70 *Cre* control, *Cre* control;B-OE, Dlx-CKO, and Dlx-CKO;B-OE brains. TorsinB overexpression prevents ChI degeneration characteristic of Dlx-CKO mice (two-way ANOVA main effect of genotype F_1,22_ = 52.45, p < 0.0001, ROSA-*Tor1b* allele F_1,22_ = 45.54, p < 0.0001, and interaction F_1,22_ = 41.90, p < 0.0001; *Cre* control n = 6, *Cre* control;B-OE n = 6, Dlx-CKO n = 7, Dlx-CKO;B-OE n = 7). (D) Representative images of P70 striatum immunostained with antibody to ChAT. ChAT+ cell density is reduced in Dlx-CKO striatum while it appears normal in Dlx-CKO;B-OE striatum.

We analyzed the brains of all four genotypes to identify potential neuropathological correlates of the B-OE-mediated behavioral rescue. TorsinB overexpression did not change cortical thickness, striatal volume, striatal Nissl+ small and medium cell number, or striatal GFAP immunoreactivity in Dlx-CKO and *Cre* control mice (Figure S6A-D). TorsinB overexpression completely prevented the loss of dorsal striatal ChIs that have been linked to motor and electrophysiologic abnormalities in dystonia mouse models [23, 29–34]. Analysis using unbiased stereology demonstrated that Dlx-CKO mice had 33.5% fewer ChIs compared to *Cre* controls, whereas the B-OE allele completely prevented this loss (Figure 4C-D). Dlx-CKO;B-OE mice did not statistically differ from *Cre* controls (mean ± SEM of 17039 ± 534 vs. 17252 ± 432 cells, respectively).

## Discussion

Our studies establish torsinB as a bidirectional modulator of torsinA dystonia-related motor phenotypes. We demonstrate that reductions of torsinB cause a dose-dependent worsening of neuropathological and motor abnormalities in multiple DYT1 models, including one containing the pathogenic ΔE disease mutation. In contrast, torsinB supplementation essentially eliminates these phenotypes, including abnormal twisting movements. These data suggest that torsinB may be an effective therapeutic target in DYT1 dystonia and are consistent with the possibility that torsinB levels play a role in determining presence or severity of symptoms in DYT1 dystonia.

These data advance understanding of the relationship of torsinB to DYT1 pathogenesis. Prior studies established a role for torsinB in the tissue selectivity and timing of torsinA-related cell biological phenotype of NE budding [17, 18], but the relationship between NE abnormalities and organismal phenotypes has not been explored previously. These new data provide proof-of-principle evidence that torsinB enhancing therapies may be disease modifying. Increasing the expression or function of torsinA itself is an alternative approach. TorsinA-targeted therapies would similarly increase levels of mutant ΔE torsinA, however, potentially worsening disease severity via a dominant negative mechanism. TorsinB-targeted therapies, in contrast, should bypass this risk. The nonphysiologic torsinB overexpression of the B-OE allele in the Nestin-Cre field caused reduced growth in Cre control mice, highlighting potential adverse consequences of this approach. The B-OE mice grossly overexpress torsinB, however, and it is likely that more modest overexpression will also provide protective effects. Future work will be needed to determine the therapeutic window of torsinB supplementation.

The interplay between torsinA LOF and torsinB levels is reminiscent of spinal muscular atrophy (SMA), a disease linked to LOF of the SMN1 gene [35]. The disease course of SMA is characterized by highly variable severity and age of onset, which is caused in large part by differences in copy number of SMN2, a duplicate of SMN1 that can attenuate SMN1 loss [36–38]. In our dystonia mouse models, a similar relationship exists where torsinB levels bidirectionally modulate the severity of torsinA LOF, worsening disease when reduced and attenuating it when overexpressed. The effect of differences in torsinB expression on DYT1 penetrance and severity in human subjects is unknown.

Neurodegeneration is one of the primary readouts in this study, whereas inherited isolated dystonia is commonly thought to be characterized by abnormal function within a structurally normal brain. Postmortem studies are limited, however, and volumetric imaging studies provide conflicting evidence [7, 39–41]. Future studies are required to fully assess potential neuropathological changes in DYT1 dystonia, especially considering that a diverse range of basal ganglia insults cause acquired dystonia [42–44]. Considerable evidence suggests that striatal cholinergic dysfunction is a key feature contributing to this disorder [29, 30, 45], supporting our focus on these cells. Our findings that torsinB overexpression prevents both striatal ChI degeneration and twisting movements further strengthens the relationship between dysfunction of these cells and DYT1-related abnormal movements.

Our findings advance the understanding of the role of torsinB in DYT1 dystonia and demonstrate that torsinB expression is a bidirectional modifier of torsinA LOF phenotypes. TorsinB overexpression is a potent disease-modifier, reducing both the prevalence and severity of abnormal movements, and fully preventing neuropathological abnormalities. These data provide a strong rationale to further explore torsinB as a target for DYT1 dystonia therapeutics.

## Methods

### Generation and maintenance of mice

*Nestin-Cre*, *Emx1-Cre*, and *Dlx5/6-Cre* were obtained from the Jackson Laboratory. *Tor1a* ΔE floxed *Tor1b* mice were generated at the University of Connecticut Center for Mouse Genome Modification. Standard methods were used to engineer the *Tor1a* and *Tor1b* loci to include the ΔE mutation and loxP sites flanking exon 3-5 of *Tor1b*. The torsinB overexpression mouse line was generated with Biocytogen using CRISPR/Extreme Genome Editing™ technology, which increases homologous recombination efficiency over standard CRISPR/Cas9 technology. Mice were maintained in our mouse colonies at the University of Michigan and the University of Texas Southwestern Medical Center. Mice were genotyped for *Tor1a*, *Cre* recombinase, *Tor1b*, and *Tor1a/Tor1b* double floxed as previously described [8, 18]. The table below contains genotyping information for the *Tor1a* ΔE floxed *Tor1b* and B-OE mouse lines:

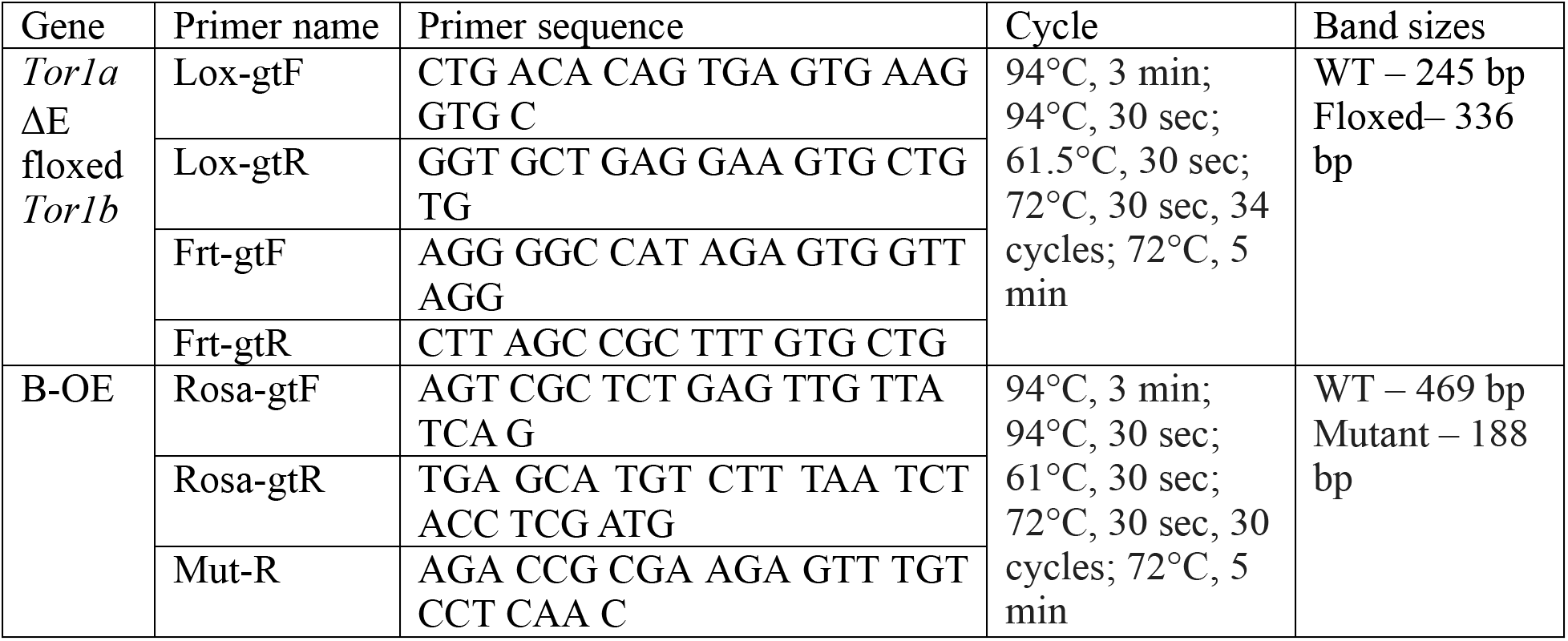

The following breeding schemes were used to generate experimental animals and littermate *Cre* controls in our studies:

TorsinB KO: *Tor1b^+/-^* x *Tor1b^+/-^*
CKO (Emx1): *Cre; Tor1a^+/-^* x *Tor1a^flx/flx^*
dCKO (Emx1): *Cre Tor1ab^+/-^* x *Tor1a/Tor1b^flx/flx^*
SKI (Emx1): *Cre*; *Tor1a^ΔE/+^* x *Tor1a^flx/flx^*
SKI-B1 (Emx1): *Cre; Tor1a^ΔE/+^* x *Tor1a/Tor1b^flx/flx^*
SKI-B0 (Emx1): *Cre; Tor1a/Tor1b^ΔE floxed torsinB/+^* x *Tor1a/Tor1b^flx/flx^*
B-OE (Dlx5/6 and Nestin): *Cre*; *Tor1a^+/-^* x *Tor1a^flx/flx^; B-OE*

Mice were housed in a temperature- and light-controlled room and provided access to food and water ad libitum. Mice of all genotypes were housed together to prevent environmental bias. Age and sex-matched littermates were used as controls for all experiments. All controls unless noted otherwise were *Cre*+ controls.

### Immunohistochemistry

Immunohistochemistry in Emx1-*Cre* studies (Figures 1–2) was performed as previously described [8]. Immunohistochemistry in torsinB rescue studies (Figures 3–4) was performed as previously described [29].

### Cell counting and morphologic analysis

CUX1 and CTIP2 immunoreactive neurons were quantified in the sensorimotor cortex as follows. A 400 μm x 1,600 μm region of interest was generated using 16 μm sagittal sections for all animals at AP bregma + 0-2 mm and ML 1.20-1.32 mm [46]. The images were viewed in ImageJ and cells expressing neuronal subtype markers were quantified by investigators blinded to genotype. Stereology on striatal Nissl stained neurons and ChAT immunoreactive neurons was performed as previously described [29]. Cortical thickness and striatal volume were measured as previously described [29]. Brain area was measured in sagittal sections by creating a contour around the brain in the section corresponding to ML 1.44 mm and measuring area of the tracing using Stereoinvestigator.

### Western Blotting

Mice were sacrificed, samples were processed, and bands were analyzed as previously described [29].

|  |  | Antibody | Company/Catalog# | Dilution |
| --- | --- | --- | --- | --- |
| IHC | Primary | Rabbit polyclonal α-GFAP | Dako #Z0334 | 1:2000 |
| | | Rabbit polyclonal α-CUX1 | Santa Cruz Biotechnology #sc-13024 | 1:200 |
| | | Rat monoclonal α-CTIP2 | Abcam #ab18465 | 1:500 |
| | | Goat polyclonal α-ChAT | Millipore AB144P | 1:200 |
| | Secondary | Donkey α-rabbit IgG, Alexa Fluor 555 conjugated | Invitrogen #A-31572 | 1:800 (GFAP),<br>1:1000 (CUX1) |
| | | Donkey α-rat IgG, Alexa Fluor 488 conjugated | Invitrogen #A-21208 | 1:1000 (CTIP2) |
| | | Donkey α-goat IgG, biotin-SP conjugated for AB144P | Jackson ImmunoResearch #705-065-147 | 1:800 (ChAT) |
| WB | Primary | Rabbit polyclonal α-torsinA | Abcam #ab34540 | 1:10,000 |
| | | Rabbit polyclonal α-torsinB | Kind gift from Schlieker lab | 1:1000 |
| | | Rabbit polyclonal α-Calnexin | Enzo #ADI-SPA-860-F | 1:20,000 |
| | Secondary | Goat α-rabbit IgG, HRP-linked | Cell Signaling #7074 | 1:20,000 (all) |

### Tail suspension testing

Mice were picked up by the tail and suspended above the home cage for one minute while and video recorded by a camera. 2 observers blinded to experimental group graded videos for duration and presence of abnormal movements. Limb clasping was defined as a sustained abnormal posture with forepaws or hind paws tightly clasped together. Trunk twisting was defined as presence of twisting of the trunk to the point where both dorsal and ventral sides of the mouse are visible. Note that trunk twisting is distinct from lateral bending that control mice exhibit in an attempt to right themselves.

### Preweaning reflexes

Forelimb hang: Mice were suspended by the forelimbs using a 3 mm wire and the latency to fall was measured at P10 age. A cutoff of 30 seconds was used.

Negative geotaxis: Mice were placed on a wire grid, and the grid was tilted to an angle of 45°with the mice facing downward. Mice were assigned a score based on the following system: 3 – no movement, 2 – able to move but unable to turn around and face upward, 1 – able to turn and face upward but unable to climb the wire grid, and 0 – able to climb the wire grid. 3 trials were conducted with 30 seconds between trials on each designed day (ages P8, P10, P12). Surface righting reflex: Mice were placed upside down on a flat surface and time to righting was measured using a stopwatch with a cutoff of 30 seconds. 3 trials were conducted on each testing day (P8, P10, P12) with 30 seconds of rest between trials.

### Statistics

All data sets are presented as mean ±SEM unless otherwise noted. Two-way ANOVAs, t-tests, and Chi square tests were performed using Graphpad prism software. Graphs were generated using the same software.

## Acknowledgements

We thank the members of the Dauer lab for their careful reading and suggestions for this manuscript. We also thank Christian Schlieker for the kind gift of the antibody to torsinB. This project was supported by: Tyler’s Hope for a Dystonia Cure Foundation; NIH R01 NS077730; Bachmann Strauss Dystonia and Parkinson Foundation.

