## Supplemental Figures for "TorsinB overexpression prevents abnormal twisting in DYT1 dystonia mouse models"

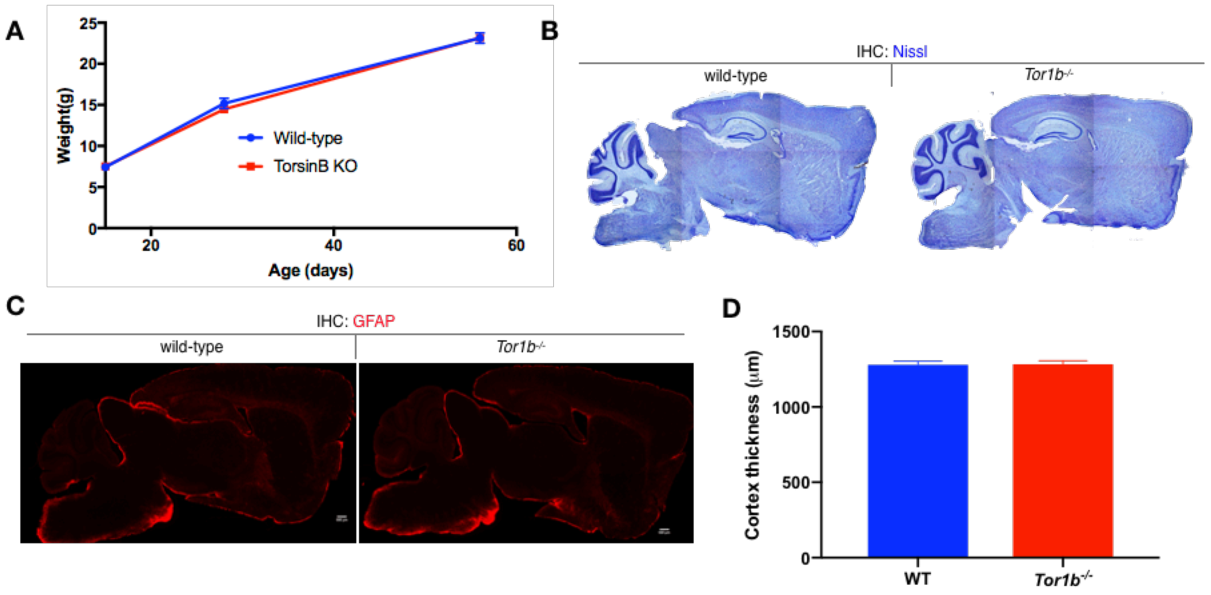


**Figure S1. TorsinB null mice exhibit no apparent organismal or neuropathological phenotype**

(A) *Tor1b*$^{-/-}$ postnatal weight gain. *Tor1b*$^{-/-}$ mutants exhibit no significant changes in body weight from age P15 to P56 compared to controls (two-way repeated measures ANOVA genotype x age, interaction $F_{2, 14} = 0.6325$, $p = 0.5458$; wild-type n = 4, torsinB KO n = 4).

(B) Nissl analysis of P56 *Tor1b*$^{-/-}$ brains. *Tor1b*$^{-/-}$ mutants did not exhibit any differences in brain morphology compared to littermate controls.

(C) GFAP staining of P56 *Tor1b*$^{-/-}$ brains. *Tor1b*$^{-/-}$ mutants did not exhibit any abnormal gliosis.

(D) Cortical thickness measurements of P56 *Tor1b*$^{-/-}$ mice. Cortical thickness does not differ significantly between WT and *Tor1b*$^{-/-}$ mice (unpaired t-test $t_6 = 0.09577$, $p = 0.9268$; wild-type n = 4, torsinB KO n = 4).

| Model | *Cre* field | Organismal/behavioral phenotype | Histologic findings | Rationale for use | Impact of torsinB change |
|---|---|---|---|---|---|
| **Emx1-CKO** | Forebrain excitatory, prominent in cortex and hippocampus | Normal appearing<br>Limb clasping in a subset of mice | Forebrain-selective neurodegeneration | Forebrain motor loop involvement<br>Mild behavioral and neuropathology findings to assess combined torsinA and B LOF | **Reduction:**<br>Worsened neuropathology and behavior |
| **Emx1-SKI** | Forebrain excitatory, prominent in cortex and hippocampus | Normal appearing<br>Limb clasping in a subset of mice | Forebrain-selective neurodegeneration milder than that see in Emx1-CKO | Forebrain motor loop involvement<br>Presence of disease mutant torsinA | **Reduction:**<br>Dose-dependent worsening of neuropathology and behavior |
| **N-CKO** | Entire nervous system | Lack of postnatal weight gain<br>Early lethality by 3rd postnatal week<br>Overtly abnormal postures at rest | Degeneration in various sensorimotor regions | Clear and robust phenotypes<br>Widespread involvement of nervous system based on the *Cre* driver | **Overexpression:**<br>Prevention of lethality, restored weight gain<br>Prevention of degeneration and gliosis |
| **Dlx-CKO** | Forebrain GABAergic and cholinergic, including all striatal neurons | Limb clasping and trunk twisting in nearly all mice that responds to drugs used in human patients | Selective degeneration of dorsal striatal cholinergic interneurons | 'Disease' model<br>Predictive validity<br>Disease course mimicking that of human patients | **Overexpression:**<br>Prevention of abnormal limb clasping and twisting movements<br>Prevention of cholinergic interneuron degeneration |

**Table S1: Features of mouse models used in this study**
Characteristics of DYT1 dystonia mouse models including extent of Cre field, behavioral phenotype, and neuropathological phenotype.

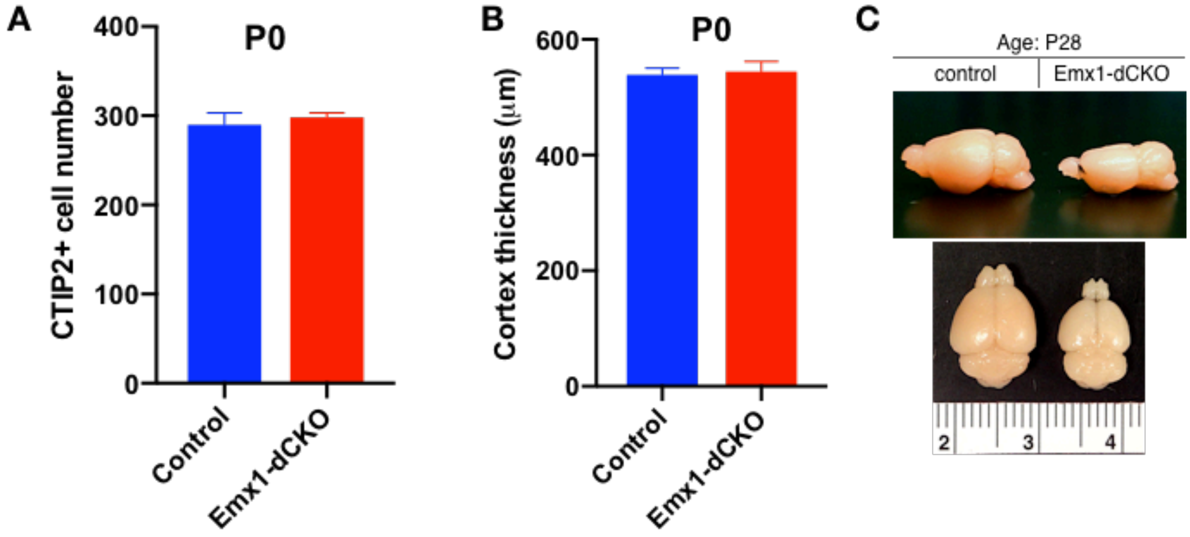


**Figure S2. Emx1-dCKO brains appear normal at birth, and gross morphological changes that emerge later are restricted to the forebrain**

(A) CTIP2 neuron counts in P0 Emx1-dCKO sensorimotor cortex. CTIP2+ neuronal counts in Emx1-dCKO mice do not differ significantly from littermate controls (non-parametric Mann-Whitney test U = 38.5, p = 0.1911; control n = 13, Emx1-dCKO n = 9).

(B) Cortical thickness measurements at P0 in Emx1-dCKO mice. Cortical thickness does differ significantly at P0 in Emx1-dCKO mutants (unpaired t-test $t_{20}$ = 0.2663, p = 0.7927; control n = 13, Emx1-dCKO n = 9).

(C) Gross morphology of Emx1-dCKO and control brains. Emx1-dCKO causes grossly apparent atrophy in the forebrain *Cre* field while hindbrain morphology appears unchanged.

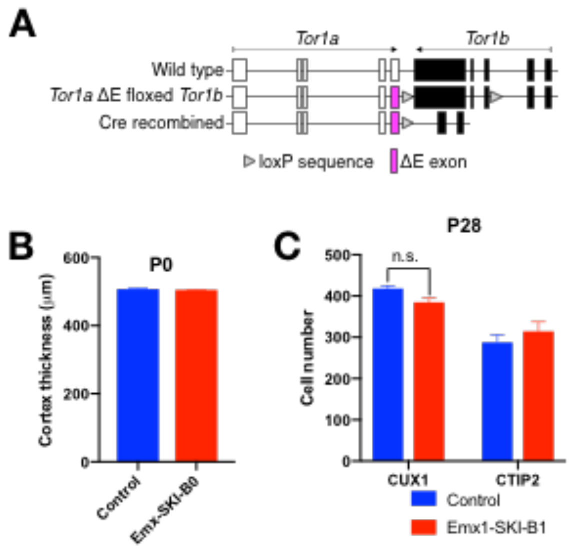


**Figure S3. *Tor1a*ΔE floxed *Tor1b* allele and Emx1-SKI-B1 cell counts**

(A) Design of the *Tor1a*ΔE floxed *Tor1b* allele.

(B) Cortical thickness of P0 Emx1-SKI-B0 mice. The thickness of the Emx1-SKI-B0 cortical tissue is unchanged at P0 (unpaired t-test $t_3 = 0.5425$, $p = 0.6252$; control n = 3, Emx1-SKI-B0 n = 2).

(C) CUX1+ and CTIP2+ counts in the sensorimotor cortex of P28 Emx1-SKI-B1 mice. There is a decrease of CUX1+ cells by 8.0% (unpaired t-test $t_4 = 2.705$, $p = 0.0538$; control n = 3, Emx1-SKI-B1 n = 5) that does not reach statistical significance in Emx1-SKI-B1 mice compared to *Cre* controls. There is no reduction of total CTIP2+ neurons in the Emx1-SKI-B1 mice compared to *Cre* controls (unpaired t-test $t_4 = 0.9128$, $p = 0.4130$; control n = 3, Emx1-SKI-B1 n = 5).

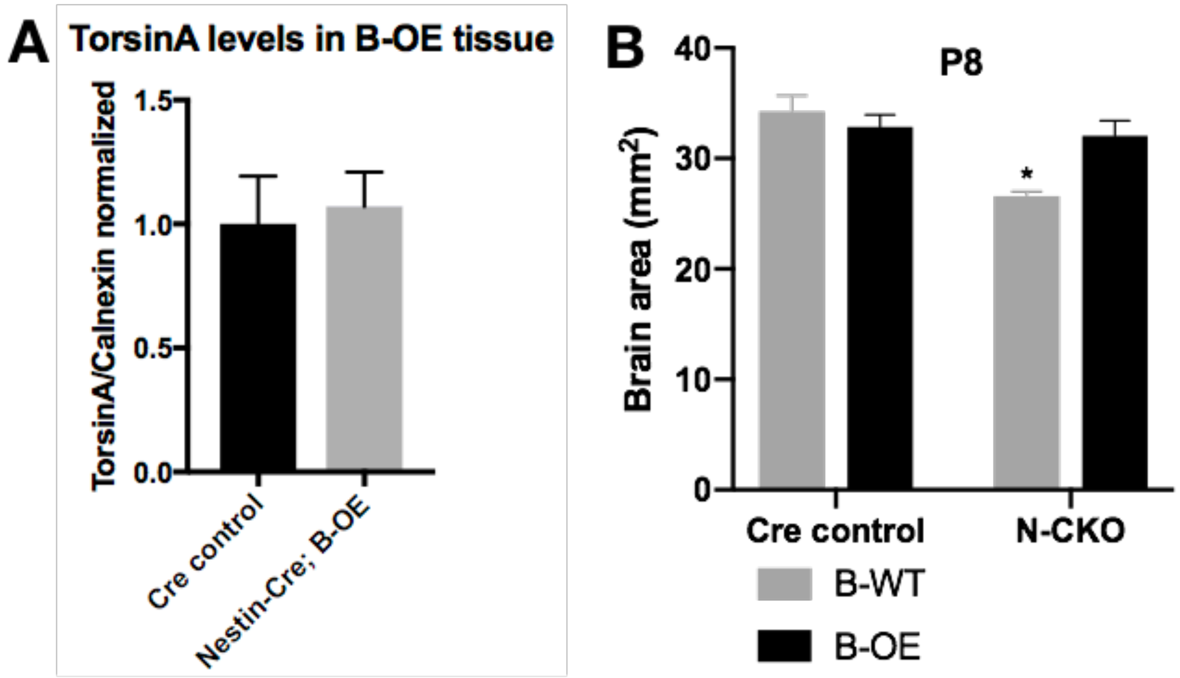


**Figure S4. Analysis of torsinA protein expression and brain size in *Nestin-Cre*;B-OE mice**

(A) Quantification of western blots of whole brain lysates from *Nestin-Cre* and *Nestin-Cre*;B-OE littermates probed with anti-torsinA antibody. TorsinB overexpression does not significantly alter levels of torsinA (unpaired t-test $t_2 = 0.2946$, $p = 0.7961$; *Nestin-Cre* n = 2, *Nestin-Cre*;B-OE n = 2).

(B) Brain area measurements of P8 *Nestin-Cre* control, *Cre* control;B-OE, N-CKO, and N-CKO;B-OE. TorsinB overexpression prevents the reduction in brain size characteristic of N-CKO mice (two-way ANOVA main effect of genotype $F_{1, 8} = 13.54$, $p = 0.0062$, main effect of torsinB level $F_{1, 8} = 3.103$, $p = 0.1162$, interaction $F_{1,8} = 8.916$, $p = 0.0174$; asterisks denote the following p-values from Tukey's multiple comparisons tests: * = $p < 0.05$; *Cre* control n = 3, *Cre* control;B-OE n = 3; N-CKO n = 3, N-CKO;B-OE n = 3).

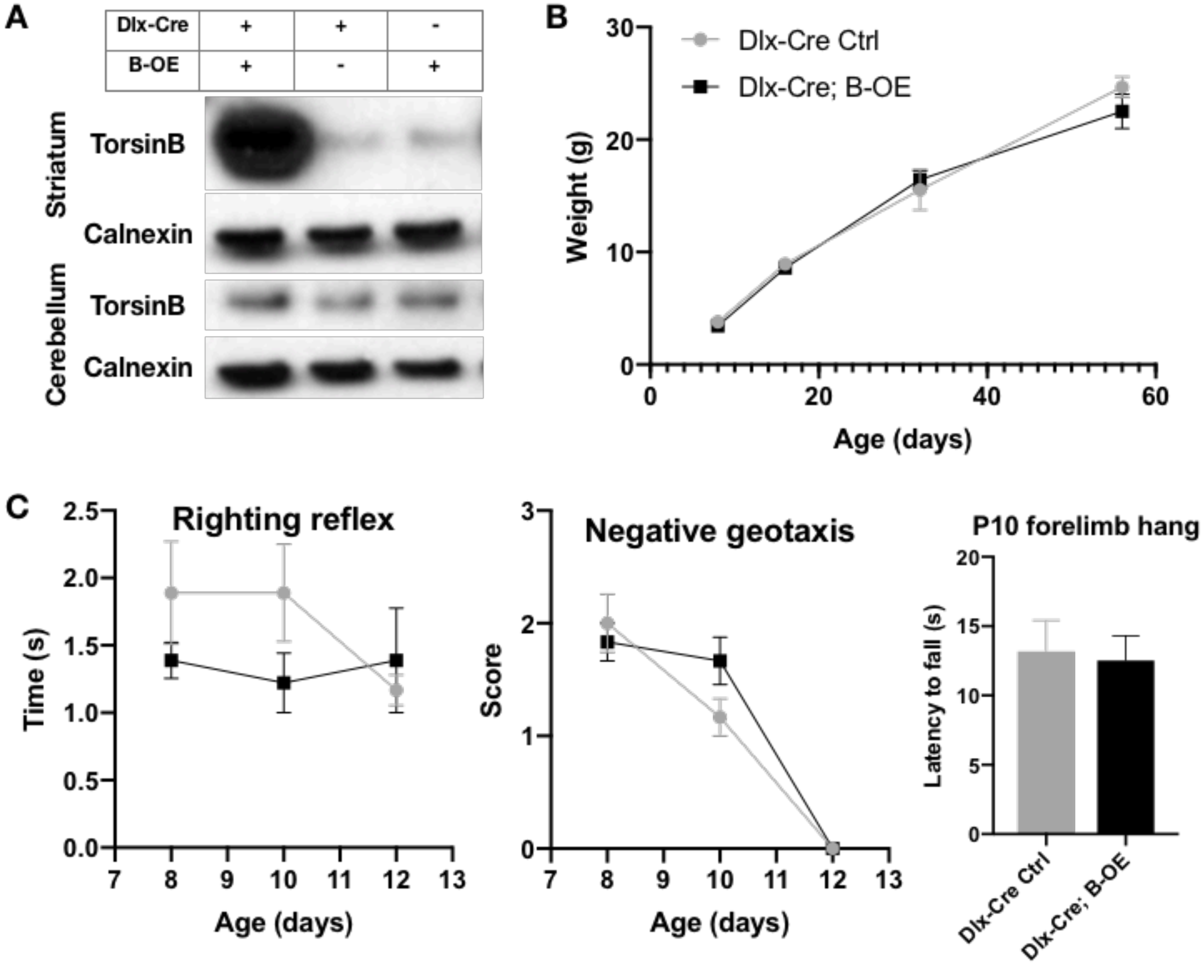


**Figure S5. TorsinB overexpression in Dlx5/6-Cre field does not impair growth or development of neonatal reflexes**

(A) Western blots of striatal and cerebellar lysates probed with anti-torsinB antibody. *Dlx5/6-Cre* selectively activates the B-OE allele in forebrain structures (striatum); torsinB expression in hindbrain structures (e.g., cerebellum) is unaffected.

(B) Postnatal weight gain of *Dlx5/6-Cre*;B-OE mice. Weigh gain is normal in *Dlx5/6-Cre*;B-OE mice compared to littermate controls (two-way repeated measures ANOVA genotype x age interaction $F_{3, 30} = 9439$, $p = 0.4318$; *Cre* control n = 6, *Cre*; B-OE n = 6).

(C) Development of neonatal reflexes in *Dlx5/6-Cre*; B-OE mice. Neonatal reflexes including righting (two-way repeated measures ANOVA genotype x age interaction $F_{2, 20} = 1.858$, $p = 0.1819$; *Cre* control n = 6, *Cre*;B-OE n = 6), negative geotaxis (two-way repeated measures ANOVA genotype x age interaction $F_{2, 20} = 2.031$, $p = 0.1574$; *Cre* control n = 6, *Cre*;B-OE n = 6), and forelimb hang (unpaired t-test $t_1 = 2317$, $p = 0.8214$; *Cre* control n = 6, *Cre*;B-OE n = 6) are normal in *Dlx5/6-Cre*;B-OE mice, indicating normal gross and motor development.

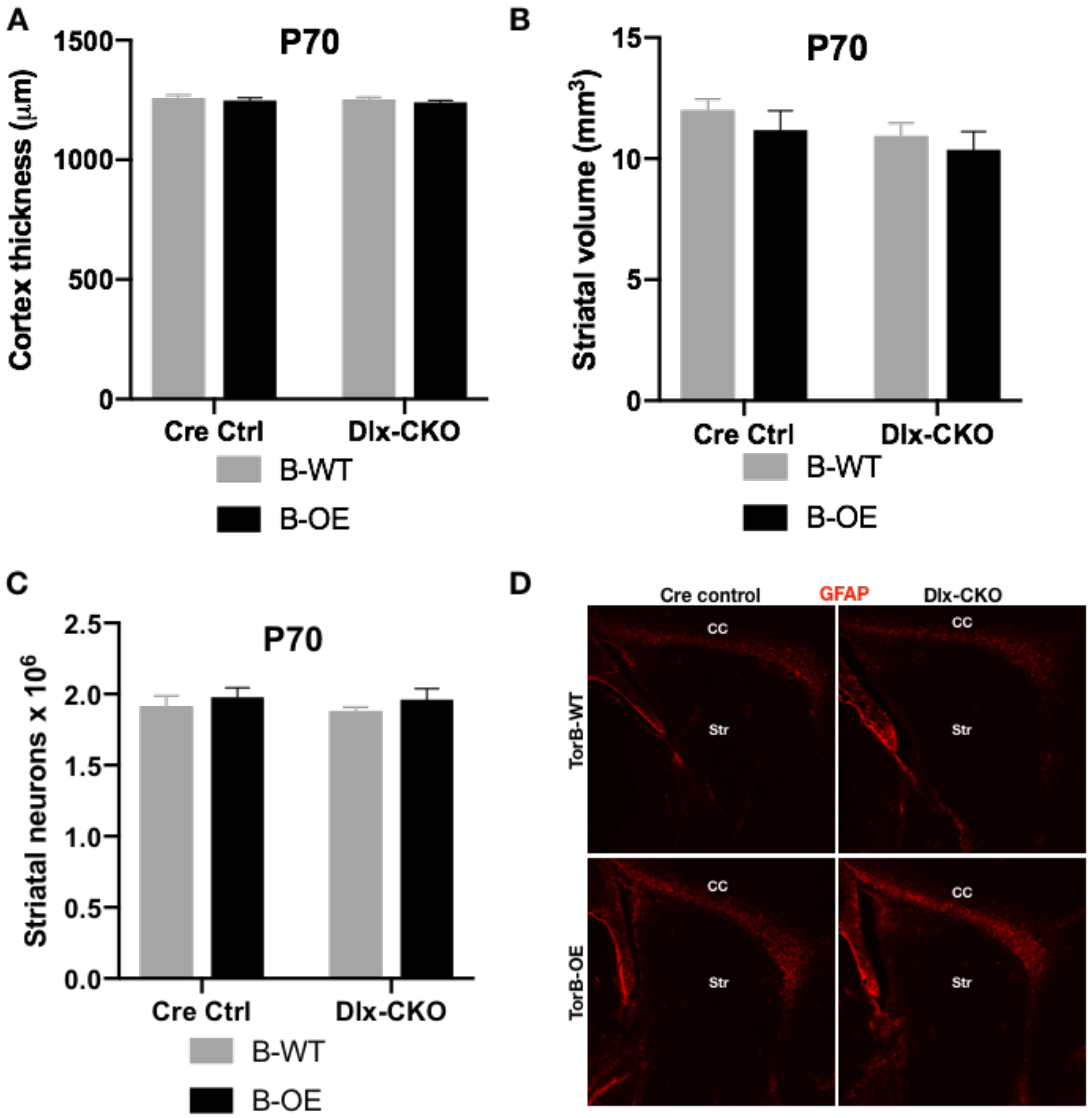


**Figure S6. TorsinB overexpression in the Dlx5/6-Cre field does not cause forebrain morphological changes or over neurodegeneration**

(A) Cortical thickness of P70 *Cre* control, *Cre* control;B-OE, Dlx-CKO, and Dlx-CKO;B-OE mice. Cortical thickness is unchanged (two-way ANOVA main genotype x torsinB level interaction $F_{1,12}$ = 0.0002485, p = 0.9877; n = 4 for all genotypes).

(B) Striatal volume of P70 *Cre* control, *Cre* control;B-OE, Dlx-CKO, and Dlx-CKO;B-OE mice. Striatal volume is unchanged (two-way ANOVA genotype x torsinB level interaction $F_{1,12}$ = 0.03657, p = 0.9515; n = 4 for all genotypes).

(C) Striatal medium and small neuron counts in P70 *Cre* control, *Cre* control;B-OE, Dlx-CKO, and Dlx-CKO;B-OE mice. The overall number of striatal neurons is unchanged two-way ANOVA genotype x torsinB level interaction $F_{1,12}$ = 0.01144, p = 0.9166; n = 4 for all genotypes).

(D) GFAP staining from *Cre* control, *Cre* control;B-OE, Dlx-CKO, and Dlx-CKO;B-OE mice. Labels show striatum (Str) and corpus callosum (CC). GFAP staining shows no reactive gliosis within the striatum of any mice.
